# Taxonomy-driven variations in snow algae color modulate albedo and energy balance in a single snow patch

**DOI:** 10.1101/2025.05.13.653574

**Authors:** Pablo Almela, James J Elser, Anthony Zmuda, Thomas Niehaus, Trinity L Hamilton

**Affiliations:** University of Minnesota Twin Cities; Flathead Lake Biological Station, University of Montana

## Abstract

In this study, we examined the reflectance, pigment composition, and community composition of three snow algae blooms showing distinct colors in the same snowfield in Glacier National Park (USA). Each color bloom was dominated by a different algae, each exhibiting a unique pigment signature but with astaxanthin as the predominant pigment across all three blooms. The spectral reflectance of red snow algae was consistently lower than that of green algae, while orange algae had intermediate reflectance values. Specifically, red algae reduced reflectance by approximately 55% across the PAR range, while green algae reduced reflectance by 25%. Red algae also demonstrated the highest radiative forcing, double that of green algae, leading to increased energy re-emission into the surrounding environment, which likely contributes to the localized melting of adjacent ice crystals. The high absorbance around 680 nm in cells with high astaxanthin content, such as the orange algae, suggests that semi-automatic detection methods could effectively identify these algae, as their spectral features remain distinct despite the presence of secondary carotenoids. Our data demonstrate the impact of snow algae taxonomic and pigment composition on the radiative balance of snowfields, underscoring taxonomy as a key determinant of bloom color under similar environmental conditions

## INTRODUCTION

The presence of light-absorbing particles (LAPs) on the surface of snow, whether abiotic (e.g., mineral dust, black carbon) or biotic (e.g., snow algae), leads to a significant decrease in albedo and an increase in melting rates (Yallop et al., 2012; Musilova et al., 2016; Stibal et al., 2017; Skiles et al., 2018; Onuma et al., 2018; Cook et al., 2020; Williamson et al., 2020; Engstrom et al., 2022). These particles often co-occur, amplifying their impact. For example, in the Alps, a combination of red snow algae and mineral dust reduced albedo by approximately 40% (Di Mauro et al., 2024). Studies focusing exclusively on snow algae have estimated albedo reductions ranging from 13% to 20% on average (Thomas and Duval, 1995; Lutz et al., 2016; Gray et al., 2020). Increased melt by biotic LAPs provide the liquid water necessary for algal growth, ensuring water availability in a highly illuminated but water-limited environment (Dial et al., 2018) and creates a feedback loop where increased algal abundance further enhances melting (Ganey et al., 2017). With permanent and seasonal snow covering up to 35% of the earth”s surface (Hell et al., 2013), clear potential exists for widespread contributions of snow algae to snow melt.

Snow algae influence the spectral albedo of snow, particularly in the visible range (400–700 nm), by darkening the surface and increasing radiative forcing (Warren and Wiscombe, 1980). Their effect on snow reflectance is closely tied to pigment composition, which dictates snow coloration. However, the relationship between environmental conditions and pigment production remains uncertain. While UV exposure and nutrient limitation are considered key regulators of secondary carotenoid synthesis, findings are inconsistent. Leya et al. (2009) observed that nitrogen starvation and high light levels in lab cultures induced carotenoid accumulation, turning algae reddish. In contrast, astaxanthin-rich algae have been reported in nitrogen-rich snow (Fujii et al., 2010), while other studies have found no direct correlation between carotenoid levels and meltwater nutrient concentrations (Müller et al., 1998). These divergent results suggest that pigment production in snow algae may not be regulated by a single set of environmental factors but rather by species-specific responses. For example, Nakashima et al. (2021) reported variations in pigment composition of colored snow associated with algal species composition. This indicates that different algal species may respond uniquely to environmental factors, which could explain variability in observed patterns of carotenoid production and impacts on albedo.

Despite growing interest in snow algae and their role in snow melt, few studies have simultaneously examined algal biomass, pigment composition, and radiative forcing while considering different snow algal colors (Gray et al., 2020; Khan et al., 2021; Halbach et al., 2024). Given that the response to environmental stressors may be species-dependent, accounting for the taxonomic identity of snow algae may be essential. To date, no study has considered the taxonomic composition of the algal community along with these factors. This gap in research limits our ability to understand and predict biological albedo reduction, and accurately use their spectral characteristics with remote sensing techniques to quantify their contribution to albedo reduction and climate feedback mechanisms.

In this study, we examined the composition, biomass abundance, pigmentation, and radiative forcing of red, orange, and green snow algae blooms occurring simultaneously in the same snowfield. The presence of different species and pigment pools under similar snow conditions allowed us to evaluate the impact of varying colors on the optical properties of an alpine snowfield. Additionally, we explored the potential of estimating algal biomass with differing carotenoid contents using spectral regions typically employed by high-resolution satellites.

## METHODS

### Field site description

Glacier National Park (GNP), referred to as Ya·qawiswit □xuki ("the place where there is a lot of ice") by the Kootenai tribe, is located in northwest Montana, United States (**Figure 1A**). During the Little Ice Age, an estimated 146 glaciers were within the current boundaries of GNP. At the end of this period, around 1850, there were about 80 glaciers in what would eventually become the national park. Only 51 of these glaciers persisted until 2005 (Martin-Mikle and Fagre, 2019). This ongoing glacial retreat makes GNP a site of great relevance for the study of climate change and cryosphere responses.

**Figure 1.**
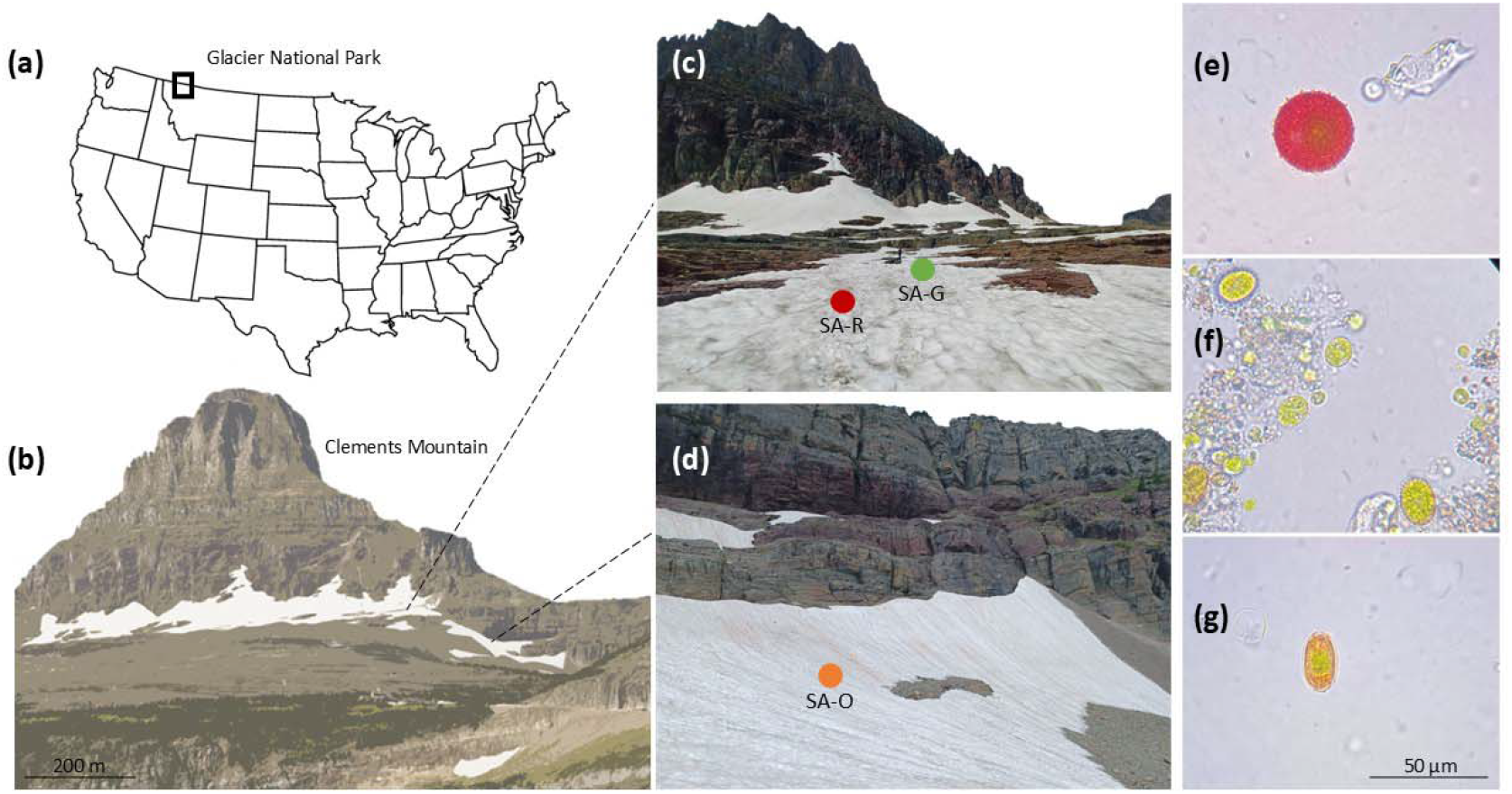
Sampling and measurement site locations. (a) Glacier National Park and (b) Clements Mountain, where red (SA-R), green (SA-G), and orange (SA-O) snow algae blooms were studied. Photos show *Sanguina nivaloides* (e), *Chloromonas alpina* (f), and *Chloromonas krienitzii* (g) fromn the different colored blooms.

Sampling was carried out on August 9 2024, on a seasonal snowfield in an open area above timber line that extended across the northeast base of Clements Mountain (48° 41” 33" N 113° 44” 10" W) at Logan Pass. The snowfield was divided into two sections by a moraine, with slopes ranging from 3° to 19°. The red and green algae blooms were found close to each other (about 10 m apart), whereas the orange snow algae bloom was in a smaller section of the snowfield, approximately 350 meters away (**Figure 1b**). Samples were visually targeted to include the three observed snowpack colors: red, orange, and green. To reduce the impact of underlying substrata on spectral albedo, data collection and snow algae sampling were conducted primarily in areas with a snowpack thickness of at least 30 cm. Control sites with snow free of visible algae (hereafter referred to as *clean snow*) were sampled at multiple locations across both sections of the snowpack to account for site-specific variability in other factors that might affect snow properties (e.g., inorganic deposition).

### Measurements of snow physical properties and spectral reflectance

In the field, and adjacent to each measurement site, snowpack depth was recorded using an avalanche probe, snow surface temperature was measured with a digital infrared thermometer, and snow water content was assessed using the SLF Snow Sensor (WSL, Switzerland) (**Supplementary Table 1**).

Prior to excavation of snow samples for laboratory analysis, spectral reflectance (as hemispherical-directional reflectance factor, HDRF) of these sites was measured using an Analytical Spectral Devices (ASD) FieldSpec® 4 hyperspectral spectroradiometer (Malvern Panalytical, USA). Although the instrument measures wavelengths from 350 to 2500 nm, this study focuses on the spectral range of 400–1300 nm, chosen for its biological relevance in capturing the light absorption and reflectance properties of pigments, while avoiding unnecessary extension into the far-infrared region where pigment signals are minimal (e.g., Di Mauro et al., 2024). Measurements were taken with a contact sensor probe that ensures consistent directional readings under uniform light intensity (halogen bulb, 2900 K color temperature), reducing contributions from stray light or temporal shifts in solar radiation. Prior to each set (i.e., color group) of snow measurements, a white reference measurement was obtained using a Spectralon panel.

To determine HDRF, surface reflectance was measured in triplicate for each of seven sampling sites for each color group (i.e., red, orange, and green) of snow algae (n = 21) (**Figure 3a,b,c**). We also measured the HDRF of *clean snow* (n = 6), to capture the variability in snowfield reflectivity outside of the red, orange, and green snow algae blooms. Reflectance (R) was determined by averaging R over two specified intervals: the visible range (400–700 nm) and the near-infrared range (700–1300 nm) (**Figure 3d,e**). In the laboratory, we measured the HDRF of algal biomass on the collection filters (see detailed methods below) to account for the reflectance of the algae while minimizing the influence of environmental factors (e.g., water content, grain size). The procedure followed was the same as in the field. A wet filter was used to measure background reflectance, which was then subtracted from the filter measurements.

To assess the potential utility of our data for remote sensing analysis, hyperspectral HDRFs were convolved with the spectral response of Sentinel 2A. Chlorophyll absorption was measured as the scaled area integral of Band 4 (665 nm) relative to Bands 3 (560 nm) and 5 (705 nm), following previously established methods (Painter et al., 2001; Gray et al., 2020; Khan et al., 2021), using the equation:

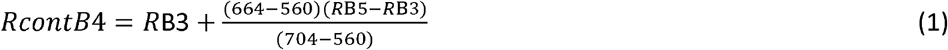

where RcontB4 is the HDRF of the continuum between Bands 3 and 5. The scaled integral of the band depth, IB4, was then calculated as

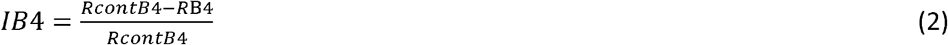

### Sample collection and biomass and inorganic particle estimations

A 2.5-cm diameter corer was used to obtain surface snow samples from the top 5 cm of each site where reflectance measurements were conducted (**Supplementary Figure 1**). The snow was placed in a sterile plastic bag and transferred to the laboratory within the next 2-3 hours of collection. After melting the sample at room temperature, a 20-µL aliquot was taken for cell counts and the remaining volume was filtered onto ashed (24 hours at 400°C) 0.7-µm pore size Whatman GF/C filters, wrapped in aluminium foil and frozen (−20 °C) until processing. Cell counts were conducted the same day of sampling by performing 9–16 replicate counts of a haemocytometer chamber (Hausser Scientific) and a light microscope (Leitz LaborLux S, with 10x objective). Photos of randomly selected cells were taken, and their lengths and widths were measured using ImageJ (Version 1.52p; National Institutes of Health, Bethesda, MD, USA). Cell volumes (μm^3^) were estimated assuming spherical shapes for red snow algae and prolate spheroid shapes for orange snow algae (**Figure 1e,g**) according to Hillebrand et al., 1999. For green snow algae, both spherical and prolate spheroid shapes were observed (**Figure 1f**).

Light-absorbing inorganic particles (LAIPs) were assessed using loss on ignition with air-dried sediments collected on pre-weighed glass fiber filters. Half of each Whatman GF/C filter was placed in a 35-mL porcelain crucible, dried at 60 °C in a convection oven for 48 hours, weighed, and then heated in a furnace at 400 °C for 24 hours. The difference in mass of the crucible and sediment before and after heating was assumed to represent the organic matter (OM) mass. The inorganic content (LAIPs) was estimated as the difference between the total mass and the OM mass.

### DNA extraction

One of the seven samples from each snow algae bloom (red, orange, green) was randomly selected to determine snow algae taxonomic composition. DNA was extracted from the filter using a DNeasy PowerSoil Kit (Qiagen, Carlsbad, CA, USA) according to the manufacturer”s instructions and Almela and Hamilton (in prep). Two negative controls were included: an extraction blank to check for kit contamination and a field blank. The field blank was an ashed 0.7-µm pore size Whatman GF/C filter washed with ultrapure (18.2 MΩ) water, and processed using the same equipment and techniques described. The concentration of DNA was determined using a Qubit DNA Assay kit (Molecular Probes, Eugene, OR, USA) and a Qubit 3.0 Fluorometer (Life Technologies, Carlsbad, CA, USA). No DNA was detected in the negative controls.

### 18S rRNA amplicon sequencing and analyses

DNA was submitted to the University of Minnesota Genomics Center (UMGC) for sequencing using a Nextera XT workflow, 2x300bp chemistry, and the primers 1391f and EukBr that target the 18S SSU rRNA (Amaral-Zettler et al., 2009; Stoeck et al., 2010).

Diversity and composition was assessed with QIIME v2-2024.10 (Bolyen et al., 2019). Briefly, cleaned and trimmed paired reads were filtered and denoised using the DADA2 plug-in (Callahan et al., 2016). For chimera identification, 340,000 training sequences were used. Identified operational taxonomic units (OTU), defined at 97% of similarity, were aligned using MAFFT (Katoh et al., 2002) and further processed to construct a phylogeny with fasttree2 (Price et al., 2010). Taxonomy was assigned to OTUs using the q2-feature-classifier (Bokulich et al., 2018) and blasted against the SILVA v138 99% 18S sequence database (Quast et al., 2012). All sequences not classified as Chlorophyta were excluded from the study. Finally, the taxonomic assignment of these sequences was confirmed by performing BLAST (Basic Local Alignment Search Tool).

### Pigment extraction and analysis

To characterize and quantify the major pigments in algal cells, half of each filter with algal biomass was placed in sterile 15-mL conical tubes, and 5 mL of a 7:2 acetone:methanol solvent was added. Cell disruption was performed by sonication (50% amplitude, 2 minutes). Samples were stored at -20 °C overnight and then sonicated again (50% amplitude, 1 minute) and centrifuged (5,000 × g, 4 °C, 10 min). An aliquot of the supernatant was filtered through a Millex® PVDF syringe filter (0.22 μm pore size). The filtered extracts were stored in glass vials at -80 °C until further analysis.

To quantify pigment concentrations, 10 μL of total extract was analyzed with an Agilent 1100 series HPLC using a Discovery® 250 × 4.6 mm C18 column (Sigma-Aldrich) with an isocratic mobile phase consisting of 50:20:30 methanol-acetonitrile-ethyl acetate flowing at 1 mL·min-1. Pigments were identified based on retention time and spectral matching to authentic standards of astaxanthin (3.17 min, 480 nm), chlorophyll-b (3.91 min, 649 nm), chlorophyll-a (4.39 min, 662 nm), and β-carotene (6.90 min, 454 nm). Diagnostic wavelengths for chlorophylls-a and -b were chosen to prevent interference with astaxanthin. The amount of compound in each peak was determined by integrating peak areas using OpenLab software (Agilent) and comparing values to those from standard curves prepared for each pigment.

Pigment measurements were calculated in volumetric units of melted snow and hence vary with the depth and volume of snow excavated from the site. From the absolute concentrations of the analysed pigments (**Figure 2b**), we calculated the pigment signature as the percentage contribution of each pigment to the total pigment composition (**Figure 2c**).

**Figure 2.**
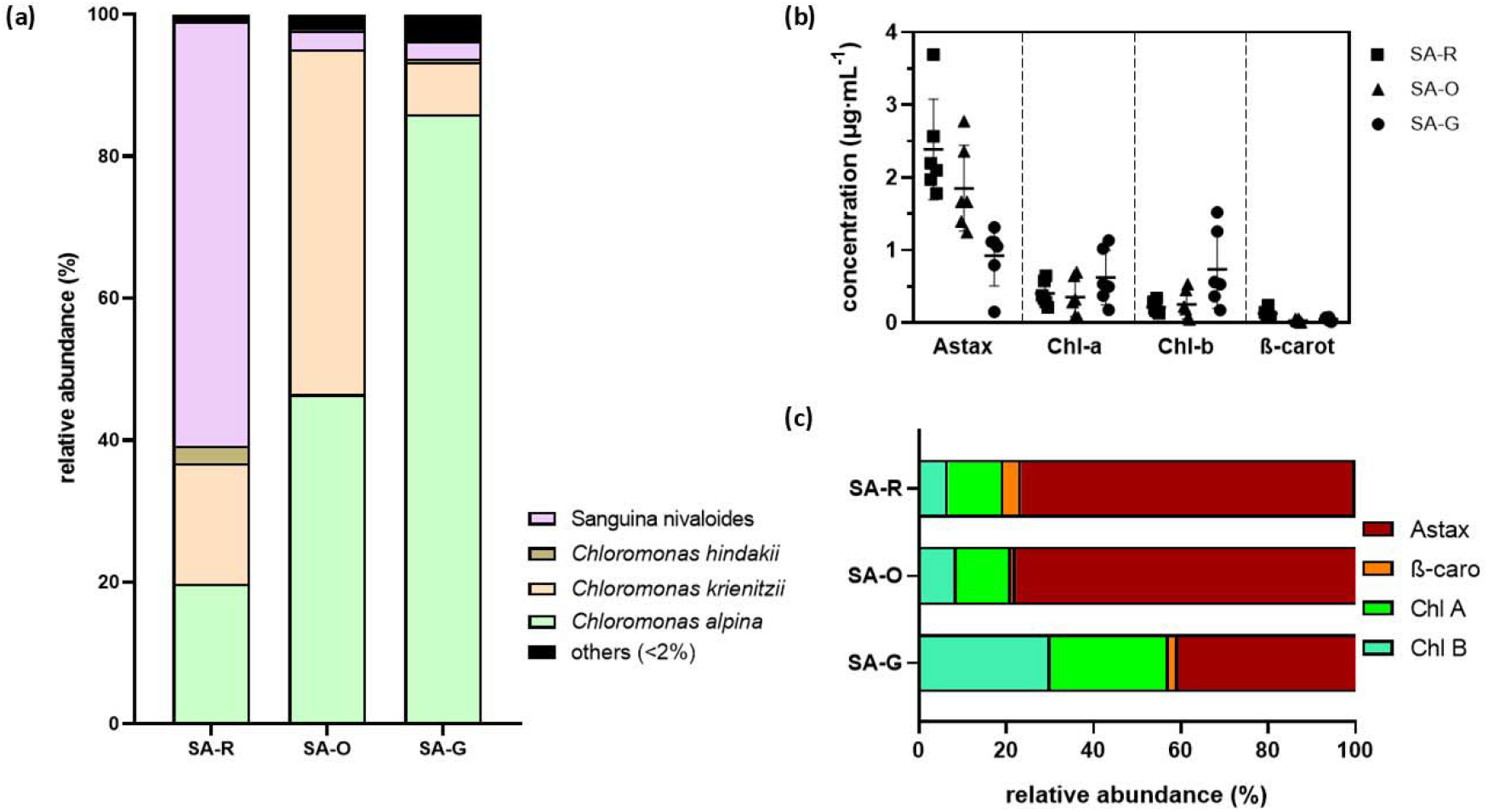
(a) Algal community composition of the three color blooms based on 18S rRNA gene metabarcoding analysis. Concentration of the pigments all-trans-Astaxanthin (Astax), Chlorophyll-a (Chl-a), Chlorophyll-b (Chl-b), and β-Carotene (β-carot) in melted snow (b), along with their relative proportions (c) for each snow algae bloom color.

**Figure 3.**
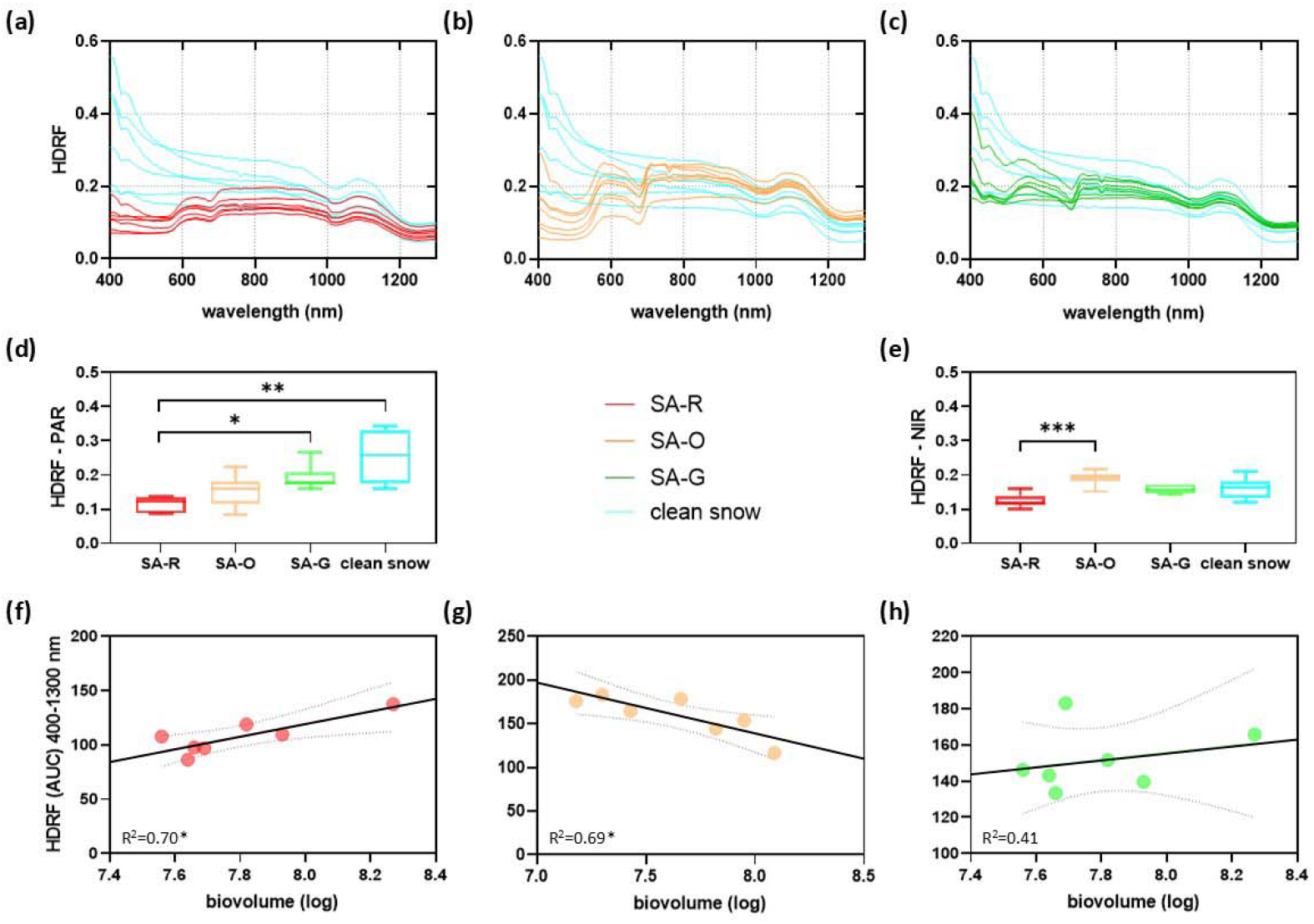
(a–c) Spectral reflectance of each snow algae type measured at Clements Mountain, GNP. Reflectance values are shown for (d) visible wavelengths (400–700 nm) and (e) near-infrared wavelengths (700–1300 nm). (f–h) Correlation between reflectance (400–1300 nm) and cell biovolume (log), with dot colors representing different snow algae types and their replicates. Asterisks (*) indicate significant differences between groups (*p < 0.05, **p < 0.01, ***p < 0.001), as determined by statistical analysis. Error bars represent the standard deviation (SD) of the mean, illustrating the variability within the samples.

### Estimate of pigment absorption

Absorptance (A(λ), defined as the ratio of absorbed to incident radiation), was calculated from reflectance (HDRF) to determine the fraction of light absorbed (i.e., not reflected) by the samples, using the specified equation:

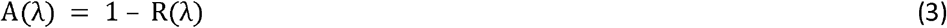

where R is the reflectance at a specific wavelength. Spectra were integrated over the PAR (400-700 nm) and NIR (700-1300 nm) regions. To estimate the photosynthetic absorptance attributed to algae in the field (bloom absorptance)(**Figure 4a,b**), we corrected for absorption relative to clean snow (Khan et al., 2021). We also estimated the absorptance attributed to the algal biomass captured on the collection filters (biomass absorptance)(**Supplementary Table 3**). Measurements of integrated intervals were presented relative to the algae biomass. This approach minimized the influence of non-pigmented algal components and other factors, such as water and impurities, ensuring consistency and comparability across measurements.

**Figure 4.**
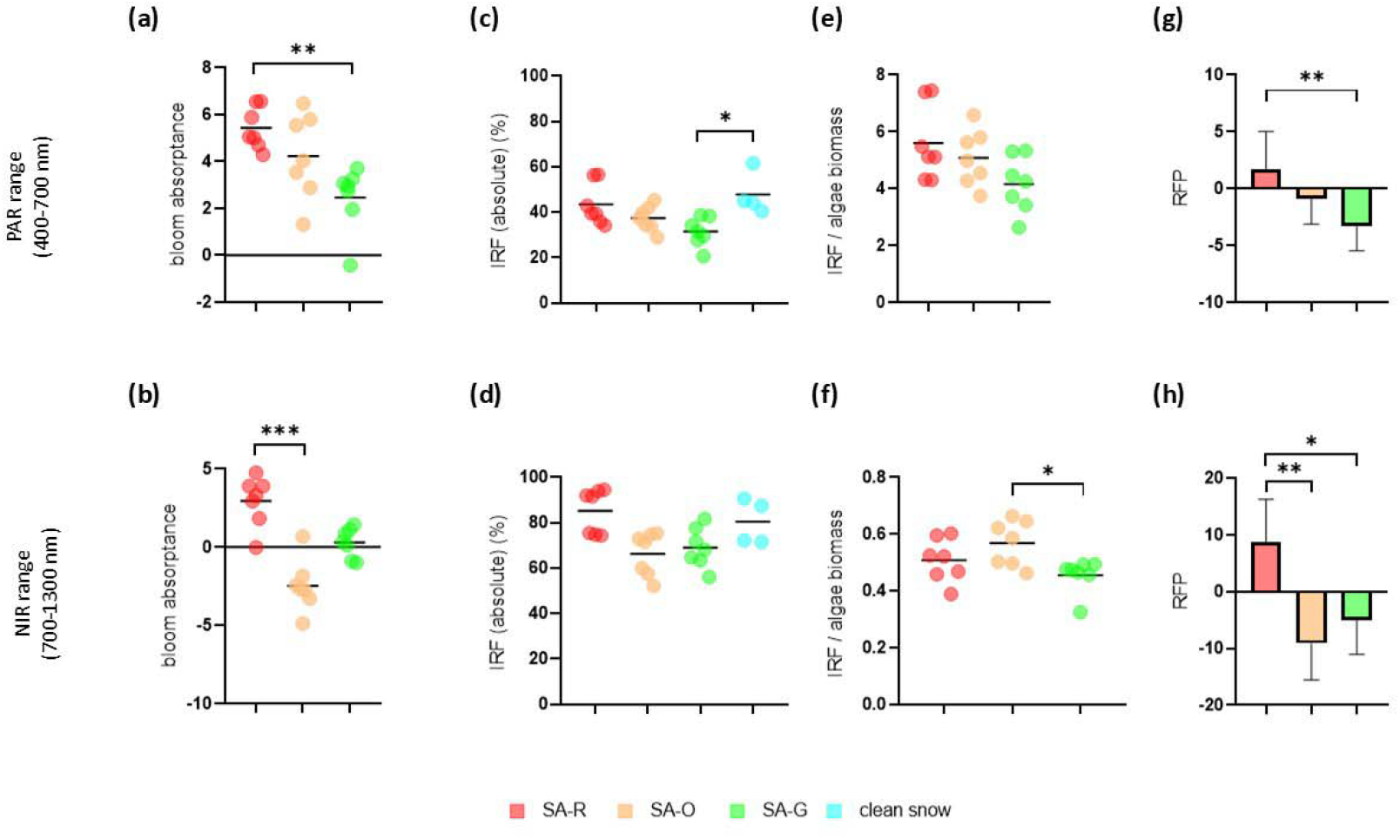
Photosynthetic absorptance attributed to algae was estimated by integrating absorption spectra over the (a) PAR (400–700 nm) and (b) NIR (700–1300 nm) regions, adjusted for absorption relative to clean snow. (c,d) Snow algal biomass absorptance was measured on filters in the lab and used to calculate (e,f) radiative forcing potential (RFP), defined as the difference between field-measured absorptance and biomass-derived absorptance from filters. Instantaneous radiative forcing (IRF) of snow algae was calculated over the (g) PAR and (h) NIR regions based on the reflectance difference between clean and algae-covered snow, accounting for solar irradiance. Asterisks (*) indicate significant differences between groups (*p < 0.05, **p < 0.01, ***p < 0.001), as determined by statistical analysis. Error bars represent the standard deviation (SD) of the mean, illustrating the variability within the samples.

### Instantaneous radiative forcing (IRF) and radiative forcing potential (RFP)

For comparison with previous studies, we calculated snow algal instantaneous radiative forcing (IRF) values over the PAR (400-700 nm) and NIR (700-1300 nm) regions following the methods used by Khan et al., 2021. IRF (W·m^−2^) was calculated based on the difference in reflectance between clean snow and algae-covered snow, and considering the solar irradiance, as described in Khan et al., 2021. The incoming spectral irradiance (W·m^−2^⍰λ_400-700_ and_700-1300_) was obtained from the PVSystems solar irradiance program (https://pvlighthouse.com.au, last access: January 2025) for the time and location of sampling. IRF was estimated as follows:

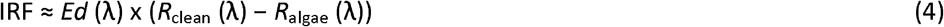

where Ed is the incoming spectral irradiance (W·m^−2^), and R is the reflectance (HDRF) for the clean or algae-covered snow (**Figure 4e,f**). Additionally, the data were presented relative to the algae biomass to standardize the measurements and facilitate comparison across the snow algae samples (**Figure 4g,h**).

We also estimated the radiative forcing potential (RFP), defined as the difference between the absorptance measured in the field samples (bloom absorptance) and the absorptance attributed to the algae biomass on the filters (biomass absorptance) (**Figure 4i,j**).

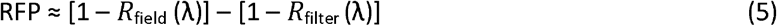

where R is the reflectance (HDRF) for the field measurement or for the corresponding filter measurements. Thus, RFP represents the proportion of the absorptance of a sample that is not attributable to its biomass and could potentially be re-emitted as heat to their surroundings.

### Data analysis and statistics

All data analysis was performed in GraphPad Prism (Version 8.0.1). To test significant differences between algal color groups, we performed a Kruskal-Wallis test, followed by Dunn”s multiple comparison test to evaluate differences in their means. We used Pearson correlations to assess relationships between biological and physical parameters of snow.

## Data Availability

Sequence Accessions—sequence data is archived at the Sequence Read Archive (SRA) at NCBI under the accessions: BioProject (PRJNA1244840) and BioSamples (SAMN47732474-SAMN47732476).

## RESULTS

### Snow characterization

Water content and temperature were generally similar across the three sites. Red patches (SA-R) had an average surface temperature of 0.2 ± 0.1 °C and a water content of 10.3 ± 1.3%.

Orange patches (SA-O) had an average surface temperature of 0.1 ± 0.0 °C and a water content of 10.1 ± 0.3%. In contrast, green patches (SA-G) had the lowest water content (8.9 ± 0.6%) and surface temperature (-0.4 ± 1.1 °C). Snow depth was significantly different between SA-R (77 ± 7.2 cm) and both SA-O (45.9 ± 11.4 cm) and SA-G (39.3 ± 12.4 cm) (**Supplementary Table 1**). For the clean snow, average surface temperature was 0.1 ± 0.0 °C, a snow depth of 70 ± 49.8 cm, and a water content of 10.4 ± 2.3%. Inorganic particle concentrations (LAIPs) on the snow varied across bloom patches (ranging from 1265.6 mg/m^2^ to 4937.6 mg/m^2^) but did not differ for different color patches (**Supplementary Table 1**).

### Algal abundance and community composition

The samples contained cells with distinct colors, morphologies and sizes. Based on these characteristics, the algal cells were classified into four types:

1. Red or deep red, spherical cells with thick cell walls (**Figure 1e**). Chloroplasts were located centrally within the cells. The mean diameter of these cells was 35.96 ± 4.9 μm.
2. Orange, oval-shaped cells with flanged cell walls (**Figure 1g**). The cells measured 25.9 ± 8.1 μm along the major axis and 14.6 ± 2.6 μm along the minor axis.
3. Green, ellipsoidal cells with smooth cell walls (**Figure 1f**). These cells were 21.1 ± 5.2 μm along the major axis and 14.7 ± 7.1 μm along the minor axis.
4. Small, green, spherical cells (**Figure 1f**). These cells had a diameter of 8.3 ± 1.3 μm. No flagellated algae were observed in any of the samples collected.

Type 1 cells were only observed in SA-R (at an average density of 3,175 cells·mL^−1^, **Supplementary Table 1**). No other cell types were observed in the red snow samples. Only Type 2 cells were found in the SA-O samples (at a density of 19,154 cells·mL^−1^). Both type 3 and type 4 cells were observed in the SA-G samples (at average densities of 99,286 and 2,286 cells·mL^−1^, respectively, corresponding to relative abundance of 97.8% and 2.2% of the total cell count). Despite the differences in cell density among colors, variations in cell size resulted in a similar total biomass per volume of melted snow across blooms (73.26 × 10^6^ vs. 32.87 × 10^6^ μm^3^·mL^−1^ for red and green blooms, respectively).

Different cell types corresponded to different taxa based on 18S rRNA analyses. *Sanguina nivaloides* dominated the snow algal community in red snow (SA-R), accounting for 60% of the total algal sequences (**Figure 2a**). In SA-G, *Chloromonas alpina* represented 86% of the sequences, while, in SA-O, *Chloromonas krienitzii* and *Chloromonas alpina* contributed 49% and 46%, respectively. Together, these three species accounted for more than 96% of the total algal sequences across the three studied blooms, albeit in different proportions.

### Pigment composition and signature

A significant positive correlation was observed between total pigment content and algal abundance in SA-R, SA-O, and SA-G, considering both cell density (R = 0.82, 0.86, and 0.90, respectively) and biovolume (R = 0.83, 0.86, and 0.90, respectively). The pigment composition varied among samples in both absolute and relative values (**Figure 2bc** and **Supplementary Table 2**). All-trans-Astaxanthin was the predominant pigment in all three samples, with the highest concentration in SA-R (2.39 ± 0.69 µg·mL^-1^), followed by SA-O (1.85 ± 0.59 µg·mL^-1^) and SA-G (0.92 ± 0.41 µg·mL-1). SA-G exhibited three-fold higher total chlorophyll (Chl-a and Chl-b) concentrations compared to SA-R and SA-O (0.62, 0.60, and 1.4 µg·mL^-1^ in SA-R, SA-O, and SA-G, respectively), whereas ß-carotene was three to four times more abundant in SA-R than in SA-O and SA-G (0.13 ± 0.06, 0.03±0.02, and 0.05±0.03 µg·mL^-1^ in SA-R, SA-O, and SA-G, respectively).

The distinct colors observed in the snow algae blooms were linked to three specific pigment signatures identified in this study, with these differences being statistically significant (2-way ANOVA, p<0.0001). SA-R was predominantly characterized by all-trans-Astaxanthin, which contributed 76.6% (±3.7) of the total pigments, and a combined total of 80.7% (±3.3%) when combined with ß-carotene. In SA-O and SA-G, the percentages of all-trans-Astaxanthin and ß-carotene combined accounted for 78.9% (±9.8) and 42.08% (±10.8) of the total pigments, respectively. In SA-G, Chl-a and Chl-b together made up 57.9% (±10.8) of the total pigments (27.4% and 30.5%, respectively). In contrast, the combined presence of Chl-a and Chl-b was lower in SA-R (19.3% ±3.3) and SA-O (21.1% ±9.8).

### Snow algae reflectance measurements

Spectral reflectance patterns were consistent with the snow and algae composition across all sites (**Figure 3abc**). Across all sites, the presence of algae caused a clear reduction in reflectance within the visible range, with distinct dips in reflectance aligning with the peak absorption regions of the dominant snow algae pigments, astaxanthin and chlorophylls. In the algae sites, red snow algae (SA-R) exhibited the lowest spectral reflectance, and green algae (SA-G) the highest (**Supplementary Table 3)**. As expected, visibly clean snow exhibited higher reflectance than the algae-containing samples, as well as high variability, especially in the visible range.

Reflectance averaged over the visible wavelength range (PAR) indicated a 55.5% decrease in SA-R (0.11±0.02) compared to the visibly clean snow (0.25±0.08), while SA-G (0.19±0.04) and SA-O (0.15±0.05) exhibited lower reductions in reflectance values (24.6% and 40.8%, respectively) (**Figure 3d**). Significant differences were observed between red and both green and clean snow samples (p < 0.05). Within the NIR wavelength range (**Figure 3e**), red snow algae (0.12 ± 0.02) reflected on average, 23.5% less energy than clean snow (0.16 ± 0.03), while green algae (0.16 ± 0.01) showed a reduction of 2.3%. Statistically significant differences were found only between red and orange snow patches (0.19 ± 0.02), with the latter reflecting less energy than clean snow.

The correlation between cell biovolume and reflectance measured in the field varied among samples. SA-R showed a significant positive correlation (R^2^ = 0.70; p = 0.02), indicating that reflectance increased with higher cell abundance (**Figure 3f**). In contrast, SA-O displayed a significant negative correlation (R^2^ = 0.69; p = 0.02), where increased algal presence corresponded to reduced reflectance (higher absorptance) (**Figure 3g**). The correlation for SA-G was positive but not statistically significant (R^2^ = 0.41; p = 0.55) (**Figure 3h**).

### Influence of snow algae blooms on snow absorptance and radiative forcing

Calculations of the snow algae bloom absorptance, based on field measurements of light reflectivity, revealed that red snow algae (SA-R) absorbed the most solar energy within the photosynthetically active radiation (PAR) range (**Supplementary Table 3**). On average, the absorptance levels of red snow algae were 120.9% higher than those of green algae and 28.6% higher than those of orange snow algae, although differences were statistically significant only in between SA-R and SA-G (**Figure 4a**). When integrating absorptance within the NIR range, red snow algae remained the most absorptive compared to snow without visible algae (*clean snow*). On average, red algae absorbed ten times more than green algae, with an even greater difference when compared to orange algae, which consistently exhibited lower absorption than clean snow (**Figure 4b**).

Absorptance measurements of algae biomass collected on filters showed a different trend than field measurements (**Supplementary Table 3**). When integrating absorptance over the PAR range and comparing it to clean snow, green snow algae absorbed 56.8% more than red algae, while orange algae absorbed 38.5% more than red algae. In the NIR range, the differences were more pronounced, although no statistically significant differences were observed.

The instantaneous radiative forcing (IRP) for clean snow in the visible range was 47.8% (±9.4) (**Figure 4e**) (**Supplementary Table 3**). Among the samples with visible snow algae, SA-R exhibited an IRP of 43.5% (±9.3), while SA-O and SA-G had lower values of 37.4% (±5.5) and 31.5% (±6.4), respectively. Compared to red algae, orange and green algae exhibited reductions in IRP of 14.1% and 27.5%, respectively, indicating a lower capacity of the cells to reflect energy back into their surroundings. In the NIR range (**Figure 4f**), red algae exhibited higher IRP values than clean snow, with averages of 85.2% (±9.7) and 80.55% (±10.0), respectively. In contrast, IRP values in SA-O and SA-G were lower: 66.3% and 69.0%, respectively, representing reductions of 22.2% and 19.0% relative to red algae. When accounting for algal biomass, the overall pattern aligned with the absolute IRF values, with red algae reflecting the most energy to their surroundings (**Figure 4g**). Orange and green algae exhibited lower IRF, with reductions of 9.2% and 25.7%, respectively, compared to red algae in the PAR range. While no significant differences were found between colors in this range, the NIR region showed significant differences between SA-O and SA-G (**Figure 4h**). In this region, orange algae had IRF values 10.5% higher than red algae and 20.0% higher than green algae.

Red snow algae exhibited the highest radiative forcing potential (RFP) among all samples, with significantly higher values than green algae in the visible range (**Figure 4g**). Notably, SA-R were the only samples to exhibit positive RFP values, indicating a greater ability to reflect a portion of the absorbed energy back to the surrounding environment. A similar trend was observed in the NIR range, where the differences were more pronounced, with significant differences between orange and green algae compared to red (**Figure 4j**). Overall, red snow algae consistently reflected a greater portion of light compared to orange and green algae.

## DISCUSSION

### Snow algae blooms of different colors impact reflectivity in different ways

The presence of algae on snow altered its optical properties in the visible part of the spectrum. Snow algae blooms can appear in a range of colors due to differences in their pigment composition, which actively influences the energy balance of snow and accelerates melting rates (Dial et al., 2018). In our study, the spectral reflectance of red (SA-R) snow algae was consistently lower than for green algae (SA-G) while orange algae (SA-O) fell in between the two types. Compared to visibly clean snow, red snow algae reduced reflectance by approximately 55% across the 400–700 nm spectral range, while green algae caused a 25% reduction, showing a significant difference and a stronger impact of red algae. NIR reflectance from 700–1300 nm was similar for red and green algae but not for orange algae, which aligns with the idea that photosynthetic pigments mainly influence the visible or PAR region without affecting NIR albedo.

To contextualize our findings, we compared them with previous studies from different geographical locations. Lutz et al. (2014) estimated a snow albedo reduction of 26% for red algae, and 31% for green algae, in the Greenland ice sheet. In the South Shetland Islands (Antarctica), Khan et al (2021) estimated contribution of 20% to reduction of snow reflectivity for red snow algae and 41% for green snow algae. Other studies, which focused solely on one algal color group, estimated a snow albedo reduction of 13–17% for red algae (Thomas and Duval, 1995; Lutz et al., 2016; Ganey et al., 2017) and around 20% for green algae blooms (Gray et al., 2020). A complicating factor in making comparisons across studies is variation in excavation depths during ground-based sampling, with many published studies using depths ranging from 5 to 10 cm. The distribution of algae at different snow depths is not yet well-documented, and pigment concentrations may vary depending on the excavation depth (Khan et al., 2021). While it is known that subsurface algae effectively absorb light (Almela et al., 2024), the full extent of this phenomenon remains unknown. However, without accounting for depth of excavation—an aspect not considered in previous studies either—our results suggest a more significant decrease in reflectance, aligning more closely with a 40% albedo reduction observed in the Alps due to the combined effect of red snow algae and mineral dust (Di Mauro et al., 2024).

We found no significant differences in the amount of inorganic material between snow algae patches differing in color or when comparing snow algae patches to clean snow. Since “clean” snow also contains LAIPs, the observed reflectance reduction can be largely attributed to the presence of algae. It is important to note that while our measurements focused on a highly localized area of a snowfield, previous studies have integrated reflectance over a larger spatial scale. As a result, our findings likely represent the potential maximum reduction in reflectance by snow algae, rather than the broad-scale albedo reduction across an entire snowfield. Under optimal conditions, however, this effect could extend more widely.

Red algae absorbed more than green algae. However, HDRF for the characteristic Chl-a absorbance centered around 680 nm (Painter et al., 2001) was significantly lower for the green algae. IB4 values derived from field-measured HDRFs ranged from 0.05 to 0.26 for SA-G and 0.01 to 0.08 for SA-R. These findings align with Gray et al., 2020 who reported that absorbance by secondary carotenoids in red algae reduces reflectance in Band 3, masking chlorophyll absorbance features within Sentinel-2 bands and making automatic detection challenging. Orange algae contain astaxanthin levels similar to those found in red algae and approximately twice those in green algae. However, IB4 values for orange algae (0.17 to 0.28), were on average higher than those for the green snow samples. This suggests that semi-automatic detection methods could effectively identify snow algae with high astaxanthin content, such as orange algae, as their spectral features remain distinct despite the presence of secondary carotenoids.

### Snow colors represented by distinct taxa and pigment compositions

In this study, we compare three snow algae blooms of distinct colors and their impact on the energy balance of an alpine snowfield. Our 18S rRNA sequencing indicates that each bloom was dominated by a different taxon. *Sanguina nivaloides* was the dominant OTU in red blooms, while *Chloromonas krienitzii* was abundant in the orange bloom. Both have previously been linked to red (Procházková et al., 2019) and orange blooms (Procházková et al., 2019), respectively. *Chloromonas alpina*, typically associated with green snow on glacial surfaces (Lutz et al., 2017), was the dominant species in the green bloom. Sequences assigned to *Chloromonas alpina* were also recovered from the orange blooms but microscopic observation revealed only one cell type in each. Snow algae are generally described as following a three-stage life cycle, characterized by changes in cell mobility, activity, pigmentation, and position within the snowpack (e.g., Hoham and Duval, 2001; Soto et al., 2023). The cycle begins with green, motile vegetative cells that emerge in spring, followed by a red, non-motile stage that remains on the snow surface and photosynthesizes during summer (Williams, Gorton and Vogelmann 2003; Remias et al., 2005). Eventually, the cells enter a dormant cyst stage. The shift from green to red cells has been linked to environmental factors such as nitrogen depletion and increased solar radiation (Bidigare et al., 1993; Remias et al., 2005; Leya et al., 2009). While this cycle is widely accepted, many of the key species involved have not been successfully cultured under laboratory conditions, indicating that current hypotheses primarily connect observed field morphologies with seasonal habitat availability (Matsumoto et al., 2024). The recovery of distinct snow algae taxa from red, orange and green blooms on the same snowpatch challenges the emphasis traditionally placed on physiological state. This finding suggests a more complex underlying mechanism and potential differences across species and cryospheric ecosystems (i.e., polar regions vs. alpine ecosystem), which may limit the direct extrapolation of results between them.

Each bloom had a distinct pigment composition; however, astaxanthin was the dominant pigment in all three communities (averaging 76.7%, 77.8%, and 40.6% in red, orange, and green snow, respectively). This emphasizes the high radiation levels experienced by cells in alpine snow, highlighting the need for protective UV adaptations. The astaxanthin:Chl-a mass ratios were higher in the red and orange snow algal samples (6.3 and 8.5, respectively) compared to the green algae (1.6 on average), indicating a higher chlorophyll proportion in the green samples. Astaxanthin likely serves as a sunscreen against harmful UV radiation, but high levels of accumulation may shade plastids, potentially reducing photosynthetic activity. Studying the pigment ratios of the algae can offer valuable ecological insights into these dynamics.

Previously documented astaxanthin:Chl-a mass ratios for snow algae differ from our findings which averaged 5.4±4.5 across all the data (red, green, and orange). For red algae, reported ratios range widely, from 56 in Halbatch et al., (2024) to 34 in Müller et al. (1998) for cells from Svalbard, with Nakashima et al. (2021) reporting values of 3 for samples from Mt. Tateyama in Japan, which are more comparable to our results. For orange blooms, Procházková et al. (2019) reported ratios between 0.04 and 0.01 in Central Europe. Studies have shown that algae from the *Sanguina* group can adjust their pigment composition in response to increasing UV radiation (Remias et al., 2005) or as populations mature during the melting season (Procházková et al., 2021). Some of the observed variability in astaxanthin:Chl-a ratios may stem from differences in analytical approaches across studies, but variations in community composition and environmental conditions could also contribute. In our samples, snow conditions (e.g., UV radiation, water content) were similar, with all three algae colors coexisting at the same location and time.

Without considering snow chemistry, which was not examined, there is no clear evidence that the three dominant species experience strong environmental constraints shaping their distribution. However, the spatial separation observed among bloom colors suggests that coexistence may be regulated by other ecological factors. A decrease in the astaxanthin:Chl-a ratio in green algae, compared to red and orange algae, may indicate increased algal depth in the snow profile, as UV radiation attenuates with depth. This aligns with our findings for green algae, which were consistently found just below the top several millimeters of the surface. Conversely, greater solar exposure may explain the significantly higher astaxanthin:Chl-a ratio in red and orange algae patches compared to green. This may indicate a photoacclimation strategy driven by a micro-habitat specialization, consistent with the higher Chl-a to Chl-b ratio observed in green algae (1:1.2) compared to red (1.9:1) and orange (1.4:1) and suggests a broader variability in pigment composition and ratios among snow algae than previously reported (e.g., Davey et al., 2019). These variations in micro-habitat occupation and pigment composition among algae colors likely optimize light capture while providing protection against excessive solar radiation, as reported in snow and ice algae from southeast Greenland glaciers (Halbach et al. 2024). However, the relationship between pigment composition and algal distribution along the vertical profile requires further investigation.

### Differences in radiative forcing among snow algae bloom colors

Red snow algae patches exhibited both the highest light absorption in the visible and near-infrared spectrum, and the greatest radiative forcing, with an average IRF that was 12% higher than that of the green patches compared to clean snow (**Figure 5**). A similar pattern was observed for RFP, where the red algae were the only samples exhibiting a significant potential to reflect absorbed energy back to the surrounding icy environment. These findings contrast with those from Antarctica (Khan et al., 2021) and the Alaskan ice field (Ganey et al., 2017), where green snow patches were shown to have a more significant impact on absorptance and IRF than red patches. These studies, along with others (e.g., Lutz et al., 2014), associate green algae with higher snow water content, which itself significantly reduces albedo (Thomas and Duval, 1995). In contrast, our results show minimal variability in snow surface properties among the samples, suggesting that differences in snow reflectivity across studies comparing green and red algae may be influenced by varying snow conditions, which could obscure the pigment-driven effects of algae. However, more research is needed to examine the changes in the vertical profile of snow”s physical properties in relation to the various algae pigments.

**Figure 5.**
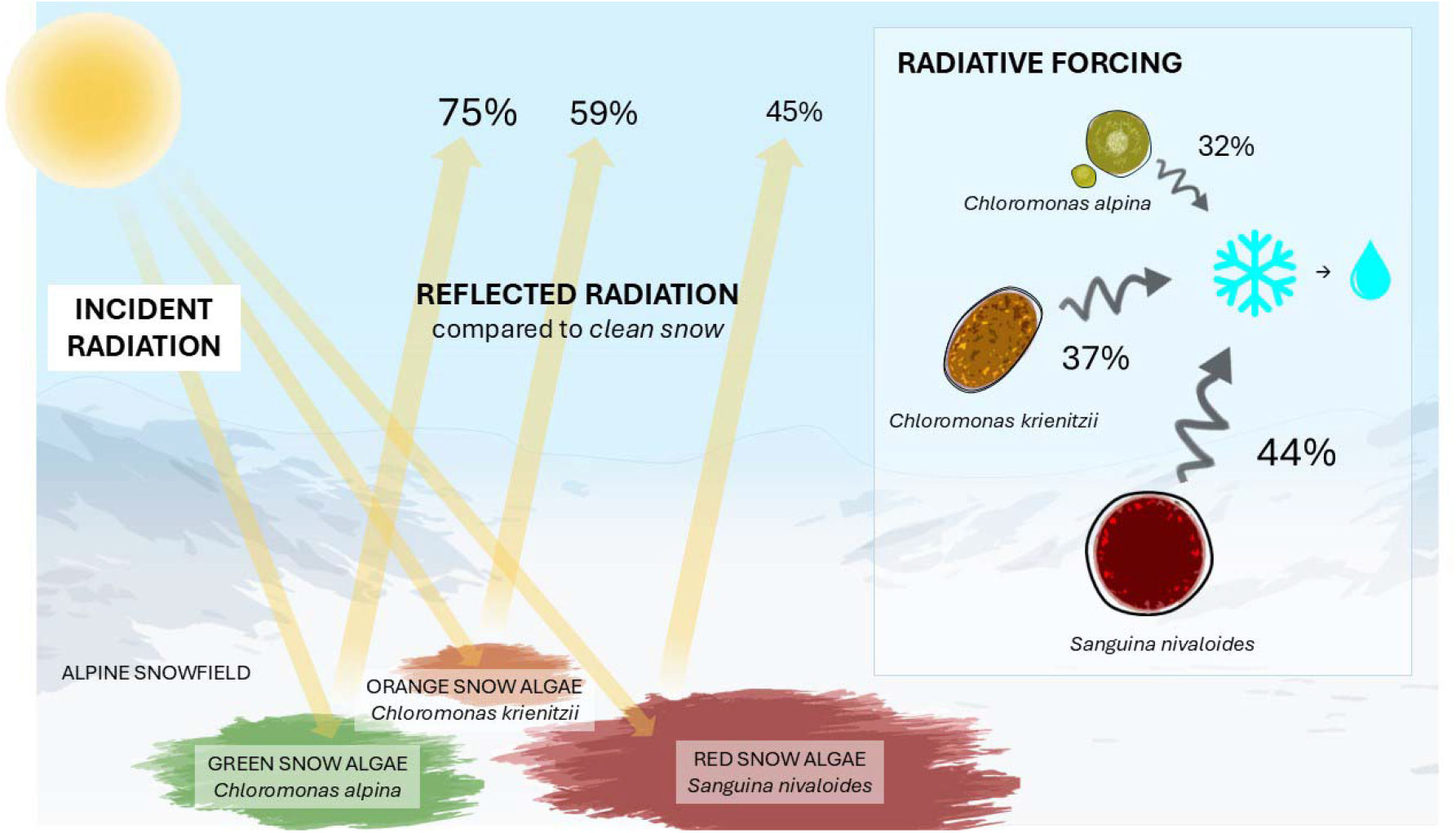
Summary of reflected radiation and radiative forcing variability in snow algae of different colors

Astaxanthin, the most abundant pigment in snow algae, occurs in various molecular forms, often bound to fatty acids and glucose. Beyond coloration, it serves as an energy storage molecule (Remias et al., 2005) and acts as a cryoprotectant by displacing water (Bidigare et al., 1993) to enhance cold survival (Řezanka et al., 2013). While these functions enhance snow algae fitness, its best characterized role is protecting thylakoid membranes from intense visible and UVA radiation, reducing photoinhibition (Remias and Lutz, 2007). This protective function also has implications for energy balance in snow algae. A proof-of-concept study identified red pigmentation as an optimal color for screening excess light and generating heat in snow algae (Dial et al., 2018), providing a selective advantage in extreme environments, where maintaining optimal cellular function is critical. Our results, some of the first studies to compare radiative forcing for different color blooms of snow algae in the alpine environment and under realistic environmental conditions, reinforce this hypothesis, demonstrating the ecological relevance of red color for snow algae.

Radiative forcing values indicated that algae with higher astaxanthin content—predominantly red and orange cells—have the greatest potential for absorbing and re-emitting energy, especially within the visible spectra. This process not only influences algal physiology but also impacts the surrounding environment, as heat absorption by pigmented cells contributes to localized melting of adjacent ice crystals (Dial et al., 2018). However, our results suggest that it is not only astaxanthin concentration that determines its impact on the snow surface”s energy balance. Correlation analyses between algal biomass—measured as biovolume to account for variations in cell size and shape—and HDRF in the field revealed that an increased presence of red algal cells on snow resulted in higher overall reflectance. However, orange algae, with a total astaxanthin content per mL of melted snow similar to red algae, exhibited an opposite correlation, absorbing energy in proportion to their biomass.

Recent findings suggest that the ability of snow algae to modify snow reflectance results from a interplay of pigments, cell density, and cell surface area, which is influenced by cell shape and size—an effect described as the Effective Albedo Reduction Surface (Almela et al., 2024). Therefore, while pigments are essential for absorbing sunlight and generating heat, an increased cell surface area can produce a similar effect even at lower pigment concentrations. However, the role of pigments in this process is more nuanced than just their abundance. The packaging of pigments can significantly influence the reflective properties of algae. A comparative study on snow and ice algae by Halbach et al. (2024) showed that the "pigment packaging effect" is especially pronounced in snow algae, with its impact on cellular energy absorption influenced by factors such as the presence of pigment-protein complexes, cell size and internal structure. This may explain the lower radiative forcing of orange algae compared to red algae in our study. However, the molecular forms of astaxanthin in snow algae have not been analyzed, and further research is needed to determine their presence and the potential effects on these differences. Moreover, despite both colors of snow algae being exposed to similar radiation at the snow surface, we have not identified a clear explanation beyond the possibility that they are distinct species that were likely sampled at different life cycle stages. These findings highlight the complex interactions between pigmentation, pigment packaging, energy balance, and environmental feedback in snow ecosystems, emphasizing the challenges of studying and predicting their response to environmental changes.

## ACKNOWLEDGEMENTS

We acknowledge the 2024 Ice and Snow Ecology class at Flathead Lake Biological Station for their discussions, which helped shape broader ideas related to this work. We sincerely thank J Joseph Giersch for providing valuable support during field sampling at Glacier National Park. This work was supported by grants #2113783 (to J.J.E.) and #2113784 (to T.L.H.) from the National Science Foundation.

## Notes

### Competing Interest Statement

The authors have declared no competing interest.

## REFERENCES

Almela, Pablo, James J Elser, J Joseph Giersch, Scott Hotaling, and Trinity L Hamilton. 2025. “Influence of Snow Cover on Albedo Reduction by Snow Algae.” mBio 16 (2): e03630–24.

Almela, Pablo, James J Elser, J Joseph Giersch, Scott Hotaling, Victoria Rebbeck, and Trinity L Hamilton. 2024. “Laboratory Experiments Suggest a Limited Impact of Increased Nitrogen Deposition on Snow Algae Blooms.” Environmental Microbiology Reports 16 (6): e70052.

Amaral-Zettler, Linda A, Elizabeth A McCliment, Hugh W Ducklow, and Susan M Huse. 2009. “A Method for Studying Protistan Diversity Using Massively Parallel Sequencing of V9 Hypervariable Regions of Small-Subunit Ribosomal RNA Genes.” PloS One 4 (7): e6372.

Bidigare, Robert R, Michael E Ondrusek, Mahlon C Kennicutt, Rodolfo Iturriaga, H Rodger Harvey, Ronald W Hoham, and Stephen A Macko. 1993. “Evidence for a Photoprotective Function for Secondary Carotenoids of Snow Algae 1.” Journal of Phycology 29 (4): 427–34.

Bokulich, NA, BD Kaehler, JR Rideout, and M Dillon. n.d. “Others (2018) Optimizing Taxonomic Classification of Markergene Amplicon Sequences with QIIME 2’s Q2-Featureclassifier Plugin.” Microbiome 6:90.

Bolyen, Evan, Jai Ram Rideout, Matthew R Dillon, Nicholas A Bokulich, Christian C Abnet, Gabriel A Al-Ghalith, Harriet Alexander, Eric J Alm, Manimozhiyan Arumugam, and Francesco Asnicar. 2019. “Reproducible, Interactive, Scalable and Extensible Microbiome Data Science Using QIIME 2.” Nature Biotechnology 37 (8): 852–57.

Callahan, Benjamin J, Paul J McMurdie, Michael J Rosen, Andrew W Han, Amy Jo A Johnson, and Susan P Holmes. 2016. “DADA2: High-Resolution Sample Inference from Illumina Amplicon Data.” Nature Methods 13 (7): 581–83.

Cook, Joseph M, Andrew J Tedstone, Christopher Williamson, Jenine McCutcheon, Andrew J Hodson, Archana Dayal, McKenzie Skiles, Stefan Hofer, Robert Bryant, and Owen McAree. 2020. “Glacier Algae Accelerate Melt Rates on the South-Western Greenland Ice Sheet.” The Cryosphere 14 (1): 309–30.

Davey, Matthew P, Louisa Norman, Peter Sterk, Maria Huete-Ortega, Freddy Bunbury, Bradford Kin Wai Loh, Sian Stockton, Lloyd S Peck, Peter Convey, and Kevin K Newsham. 2019. “Snow Algae Communities in Antarctica: Metabolic and Taxonomic Composition.” New Phytologist 222 (3): 1242–55.

Di Mauro, B, R Garzonio, C Ravasio, V Orlandi, G Baccolo, S Gilardoni, D Remias, B Leoni, M Rossini, and R Colombo. 2024. “Combined Effect of Algae and Dust on Snow Spectral and Broadband Albedo.” Journal of Quantitative Spectroscopy and Radiative Transfer 316:108906.

Dial, Roman J, Gerard Q Ganey, and S McKenzie Skiles. 2018. “What Color Should Glacier Algae Be? An Ecological Role for Red Carbon in the Cryosphere.” FEMS Microbiology Ecology 94 (3): fiy007.

Engstrom, Casey B, Scott N Williamson, John A Gamon, and Lynne M Quarmby. 2022. “Seasonal Development and Radiative Forcing of Red Snow Algal Blooms on Two Glaciers in British Columbia, Canada, Summer 2020.” Remote Sensing of Environment 280:113164.

Fujii, Masanori, Yoshinori Takano, Hisaya Kojima, Tamotsu Hoshino, Ryouichi Tanaka, and Manabu Fukui. 2010. “Microbial Community Structure, Pigment Composition, and Nitrogen Source of Red Snow in Antarctica.” Microbial Ecology 59:466–75.

Ganey, Gerard Q, Michael G Loso, Annie Bryant Burgess, and Roman J Dial. 2017. “The Role of Microbes in Snowmelt and Radiative Forcing on an Alaskan Icefield.” Nature Geoscience 10 (10): 754–59.

Gray, Andrew, Monika Krolikowski, Peter Fretwell, Peter Convey, Lloyd S Peck, Monika Mendelova, Alison G Smith, and Matthew P Davey. 2020. “Remote Sensing Reveals Antarctic Green Snow Algae as Important Terrestrial Carbon Sink.” Nature Communications 11 (1): 2527.

Halbach, Laura, Katharina Kitzinger, Martin Hansen, Sten Littmann, Liane G Benning, James A Bradley, Martin J Whitehouse, Malin Olofsson, Rey Mourot, and Martyn Tranter. 2025. “Single-Cell Imaging Reveals Efficient Nutrient Uptake and Growth of Microalgae Darkening the Greenland Ice Sheet.” Nature Communications 16 (1): 1521.

Hell, Katherina, Arwyn Edwards, Jakub Zarsky, Sabine M Podmirseg, Susan Girdwood, Justin A Pachebat, Heribert Insam, and Birgit Sattler. 2013. “The Dynamic Bacterial Communities of a Melting High Arctic Glacier Snowpack.” The ISME Journal 7 (9): 1814–26.

Hillebrand, Helmut, Claus-Dieter Dürselen, David Kirschtel, Utsa Pollingher, and Tamar Zohary. 1999. “Biovolume Calculation for Pelagic and Benthic Microalgae.” Journal of Phycology 35 (2): 403–24.

Jones, H Gerald. 2001. Snow Ecology: An Interdisciplinary Examination of Snow-Covered Ecosystems. Cambridge University Press.

Katoh, Kazutaka, Kazuharu Misawa, Kei-ichi Kuma, and Takashi Miyata. 2002. “MAFFT: A Novel Method for Rapid Multiple Sequence Alignment Based on Fast Fourier Transform.” Nucleic Acids Research 30 (14): 3059–66.

Khan, Alia L, Heidi M Dierssen, Ted A Scambos, Juan Höfer, and Raul R Cordero. 2021. “Spectral Characterization, Radiative Forcing and Pigment Content of Coastal Antarctic Snow Algae: Approaches to Spectrally Discriminate Red and Green Communities and Their Impact on Snowmelt.” The Cryosphere 15 (1): 133–48.

Leya, Thomas, Andreas Rahn, Cornelius Lütz, and Daniel Remias. 2009. “Response of Arctic Snow and Permafrost Algae to High Light and Nitrogen Stress by Changes in Pigment Composition and Applied Aspects for Biotechnology.” FEMS Microbiology Ecology 67 (3): 432– 43.

Lutz, Stefanie, Alexandre M Anesio, Arwyn Edwards, and Liane G Benning. 2017. “Linking Microbial Diversity and Functionality of Arctic Glacial Surface Habitats.” Environmental Microbiology 19 (2): 551–65.

Lutz, Stefanie, Alexandre M Anesio, Susana E Jorge Villar, and Liane G Benning. 2014. “Variations of Algal Communities Cause Darkening of a Greenland Glacier.” FEMS Microbiology Ecology 89 (2): 402–14.

Lutz, Stefanie, Alexandre M Anesio, Rob Raiswell, Arwyn Edwards, Rob J Newton, Fiona Gill, and Liane G Benning. 2016. “The Biogeography of Red Snow Microbiomes and Their Role in Melting Arctic Glaciers.” Nature Communications 7 (1): 11968.

Martin-Mikle, Chelsea J, and Daniel B Fagre. 2019. “Glacier Recession since the Little Ice Age: Implications for Water Storage in a Rocky Mountain Landscape.” Arctic, Antarctic, and Alpine Research 51 (1): 280–89.

Matsumoto, Maya, Clare Hanneman, A_G Camara, Stacy A Krueger-Hadfield, Trinity L Hamilton, and Robin B Kodner. 2024. “Hypothesized Life Cycle of the Snow Algae Chlainomonas Sp.(Chlamydomonadales, Chlorophyta) from the Cascade Mountains, USA.” Journal of Phycology 60 (3): 724–40.

Müller, T, W Bleiß, C-D Martin, S Rogaschewski, and G Fuhr. 1998. “Snow Algae from Northwest Svalbard: Their Identification, Distribution, Pigment and Nutrient Content.” Polar Biology 20:14–32.

Musilova, Michaela, Martyn Tranter, Jonathan L Bamber, Nozomu Takeuchi, and Alexandre MB Anesio. 2016. “Experimental Evidence That Microbial Activity Lowers the Albedo of Glaciers.” Geochemical Perspectives Letters 2 (2): 105–16.

Nakashima, Tomomi, Jun Uetake, Takahiro Segawa, Lenka Procházková, Akane Tsushima, and Nozomu Takeuchi. 2021. “Spatial and Temporal Variations in Pigment and Species Compositions of Snow Algae on Mt. Tateyama in Toyama Prefecture, Japan.” Frontiers in Plant Science 12:689119.

Onuma, Yukihiko, Nozomu Takeuchi, Sota Tanaka, Naoko Nagatsuka, Masashi Niwano, and Teruo Aoki. 2018. “Observations and Modelling of Algal Growth on a Snowpack in North-Western Greenland.” The Cryosphere 12 (6): 2147–58.

Painter, Thomas H, Brian Duval, William H Thomas, Maria Mendez, Sara Heintzelman, and Jeff Dozier. 2001. “Detection and Quantification of Snow Algae with an Airborne Imaging Spectrometer.” Applied and Environmental Microbiology 67 (11): 5267–72.

Price, Morgan N, Paramvir S Dehal, and Adam P Arkin. 2010. “FastTree 2–Approximately Maximum-Likelihood Trees for Large Alignments.” PloS One 5 (3): e9490.

Prochazkova, Lenka, Thomas Leya, Heda Křížková, and Linda Nedbalova. 2019. “Sanguina Nivaloides and Sanguina Aurantia Gen. et Spp. Nov.(Chlorophyta): The Taxonomy, Phylogeny, Biogeography and Ecology of Two Newly Recognised Algae Causing Red and Orange Snow.” FEMS Microbiology Ecology 95 (6): fiz064.

Procházková, Lenka, Daniel Remias, Wolfgang Bilger, Heda Křížková, Tomáš Řezanka, and Linda Nedbalová. 2020. “Cysts of the Snow Alga Chloromonas Krienitzii (Chlorophyceae) Show Increased Tolerance to Ultraviolet Radiation and Elevated Visible Light.” Frontiers in Plant Science 11:617250.

Procházková, Lenka, Daniel Remias, Andreas Holzinger, Tomáš Řezanka, and Linda Nedbalová. 2021. “Ecophysiological and Ultrastructural Characterisation of the Circumpolar Orange Snow Alga Sanguina Aurantia Compared to the Cosmopolitan Red Snow Alga Sanguina Nivaloides (Chlorophyta).” Polar Biology 44 (1): 105–17.

Quast, Christian, Elmar Pruesse, Pelin Yilmaz, Jan Gerken, Timmy Schweer, Pablo Yarza, Jörg Peplies, and Frank Oliver Glöckner. 2012. “The SILVA Ribosomal RNA Gene Database Project: Improved Data Processing and Web-Based Tools.” Nucleic Acids Research 41 (D1): D590–96.

Remias, DANIEL, and CORNELIUS Lütz. 2007. “Characterisation of Esterified Secondary Carotenoids and of Their Isomers in Green Algae: A HPLC Approach.” Algological Studies 124:85–94.

Remias, Daniel, Ursula Lütz-Meindl, and Cornelius Lütz. 2005. “Photosynthesis, Pigments and Ultrastructure of the Alpine Snow Alga Chlamydomonas Nivalis.” European Journal of Phycology 40 (3): 259–68.

Soto, Daniela F, Iván Gómez, and Pirjo Huovinen. 2023. “Antarctic Snow Algae: Unraveling the Processes Underlying Microbial Community Assembly during Blooms Formation.” Microbiome 11 (1): 200.

Stibal, Marek, Jason E Box, Karen A Cameron, Peter L Langen, Marian L Yallop, Ruth H Mottram, Alia L Khan, Noah P Molotch, Nathan AM Chrismas, and Filippo Calì Quaglia. 2017. “Algae Drive Enhanced Darkening of Bare Ice on the Greenland Ice Sheet.” Geophysical Research Letters 44 (22): 11–463.

Stoeck, Thorsten, David Bass, Markus Nebel, Richard Christen, Meredith DM Jones, Hans-Werner Breiner, and Thomas A Richards. 2010. “Multiple Marker Parallel Tag Environmental DNA Sequencing Reveals a Highly Complex Eukaryotic Community in Marine Anoxic Water.” Molecular Ecology 19:21–31.

Thomas, William H, and Brian Duval. 1995. “Sierra Nevada, California, USA, Snow Algae: Snow Albedo Changes, Algal-Bacterial Interrelationships, and Ultraviolet Radiation Effects.” Arctic and Alpine Research 27 (4): 389–99.

Williams, William E, Holly L Gorton, and Thomas C Vogelmann. 2003. “Surface Gas-Exchange Processes of Snow Algae.” Proceedings of the National Academy of Sciences 100 (2): 562–66.

Williamson, Christopher J, Karen A Cameron, Joseph M Cook, Jakub D Zarsky, Marek Stibal, and Arwyn Edwards. 2019. “Glacier Algae: A Dark Past and a Darker Future.” Frontiers in Microbiology 10:524.

Wiscombe, Warren J, and Stephen G Warren. 1980. “A Model for the Spectral Albedo of Snow. I: Pure Snow.” Journal of Atmospheric Sciences 37 (12): 2712–33.

Yallop, Marian L, Alexandre M Anesio, Rupert G Perkins, Joseph Cook, Jon Telling, Daniel Fagan, James MacFarlane, Marek Stibal, Gary Barker, and Chris Bellas. 2012. “Photophysiology and Albedo-Changing Potential of the Ice Algal Community on the Surface of the Greenland Ice Sheet.” The ISME Journal 6 (12): 2302–13.

